# Anuran assemblage changes along small-scale phytophysiognomies in natural Brazilian grasslands

**DOI:** 10.1101/2020.07.31.229310

**Authors:** Diego Anderson Dalmolin, Volnei Mathies Filho, Alexandro Marques Tozetti

**Affiliations:** Laboratório de Metacomunidades, Departamento de Ecologia, Universidade Federal do Rio Grande do Sul, Porto Alegre, Brazil; Fundação Universidade Federal do Rio Grande, Rio Grande, Rio Grande do Sul, Brasil; Laboratório de Ecologia de Vertebrados Terrestres, Universidade do Vale do Rio dos Sinos, Avenida Unisinos 950, 93022-000 São Leopoldo, Rio Grande do Sul, Brazil

**Keywords:** Dissimilarity, anuran, south Brazil, Taim, forest, grasslands

## Abstract

We studied the species composition of frogs in two phytophysiognomies (grassland and forest) of a Ramsar site in southern Brazil. We aimed to assess the distribution of species on a small spatial scale and dissimilarities in community composition between grassland and forest habitats. The sampling of individuals was carried out through pitfall traps and active search in the areas around the traps. We evaluated the existence of these differences by using permutational multivariate analysis of variance and multivariate dispersion. We found 13 species belonging to six families. Leptodactylidae and Hylidae were the most representative families. The compositional dissimilarity was higher between the sampling sites from different phytophysiognomies than within the same phytophysiognomy, suggesting that forest and grassland drive anuran species composition differently. Also, the difference in anuran species composition between the sampling sites within the forest was considerably high. Based on our results, we could assume that the phytophysiognomies evaluated here offer quite different colonization opportunities for anurans, especially those related to microhabitat characteristics, such as microclimate variables.

## Introduction

The composition of species in communities is a result of a complex interaction between organisms and environmental characteristics. The presence of a singular species in an assemblage depends on intrinsic (such as the individual’s physiological needs for homeostatic maintenance) and extrinsic processes (such as competition, predation and other biotic interactions). Combined, both of them determine the possibility of a given species to colonize a habitat, as well their long-term maintenance in there (Schluter & Ricklefs 1993; Ximenez et al. 2014). It is assumed that heterogeneous habitats comprise a richer biota (Semlitsch et al. 2015) since they offer many different microhabitats, which precisely favors the establishment of species without high niche overlap (Barrows & Allen 2010; Oliveira et al. 2013)

The concept of habitat heterogeneity is related to much confusion, being frequently used as a synonym of habitat complexity. The process of describing and evaluating the habitat structure is one of the challenges for carrying out ecological studies (Stein & Kreft 2015). In structural terms, environments can be described by (i) their complexity - vertical development of vegetation - and (ii) their heterogeneity - horizontal variation in descriptors such as density, dominance and frequency of plant species that compose them (August 1983). In a review on the topic, Tews et al. (2004) defend the idea that the concept of habitat heterogeneity includes both vertical and horizontal structure of plant community - a concept adopted in the present study. The possibility of assessing the effect of habitat heterogeneity on the structuring of their communities comes up against the huge number of areas never sampled in the Neotropical region (Garcia & Vinciprova 2003). This fact also has implications for the establishment of conservation strategies for potentially threatened species - as is the case with many Brazilian anurans (Silvano & Segalla 2005).

The variety of available habitats and the high levels of species endemism throughout the Brazilian biomes makes it an adequate location to assess the role of environmental heterogeneity in the activity patterns and community structure of anurans (Silvano & Segalla 2005). However, most studies on anuran ecology in Brazil are concentrated in forest environments, especially in the Cerrado and the Atlantic Forest biomes (da Silva et al. 2011a; Iop et al. 2012; Prado & Rossa-Feres, 2014; Saccol et al., 2017). The Pampa grassland formations and the coastal portion of southern Brazil are home to anuran species that are common in open habitats (Núñez et al. 2004, Vasconcelos et al. 2019; Dalmolin et al. 2019). Although the landscape of these environments consists predominantly of open areas (Machado et al. 2012), some forest patches can also be observed (such as swamp forests and dry or sandy forests).

Studies that describe changes in amphibian assemblages over structural changes in vegetation cover can contribute to propose scenarios of landscape changes (Da Silva et al. 2011a). The process of forests expanding over grasslands under climate change or even converting forested areas into human use would reflect changes in the amphibian composition (Manenti et al. 2013; Oliveira & Rocha 2015). In this sense, our work aimed to: (i) verify the distribution of anuran species on a small spatial scale; (ii) describe the association between the patterns of dissimilarity in anuran composition between communities of different phytophysiognomies (forest and grassland).

The study design on a small spatial scale is interesting to assess adjacent habitats whose probability of colonization is theoretically the same for all species in the region (Ximenez et al. 2014). In addition, the habitat elements linked to the micro-habitat scale are important descriptors of the spatial partition of species (Freitas et al. 2002, Oliveira & Rocha 2015) in particular because they deal with transition zones in a refined way (Bersier et al. 2007). The detection of differences in anuran assemblages between habitats provides a strong indication of the segregation process of these species along a vegetation gradient (Da Silva et al. 2011a; Prado & Rossa-Feres 2014). We expected to find greater differences in composition in places belonging to different phytophysiognomies than between areas belonging to the same phytophysiognomies, since the environmental and microenvironmental conditions must be similar in these areas, thus reducing their composition dissimilarity (Da Silva et al. 2011).

## Material and Methods

### Study site

The study was carried out at the Estação Ecológica do Taim (ESEC Taim; S 32°32’16.6 “W 052°32’16.7”), located in the municipality of Rio Grande, Brazil’s extreme south (Figure 1). ESEC Taim encompasses ecosystems of great relevance and, since 2017, was recognized as a Ramsar site (Ramsar Wetland of International Importance; Ramsar, 2018). The landscape consists of wetlands and grasslands in association with remnants of “restinga” and flooded forests, which are dominated by cork trees (*Erythrina crista-galli*) and fig trees (*Ficus organensis*; Waechter & Jarenkow 1998).

**Figure 1:**
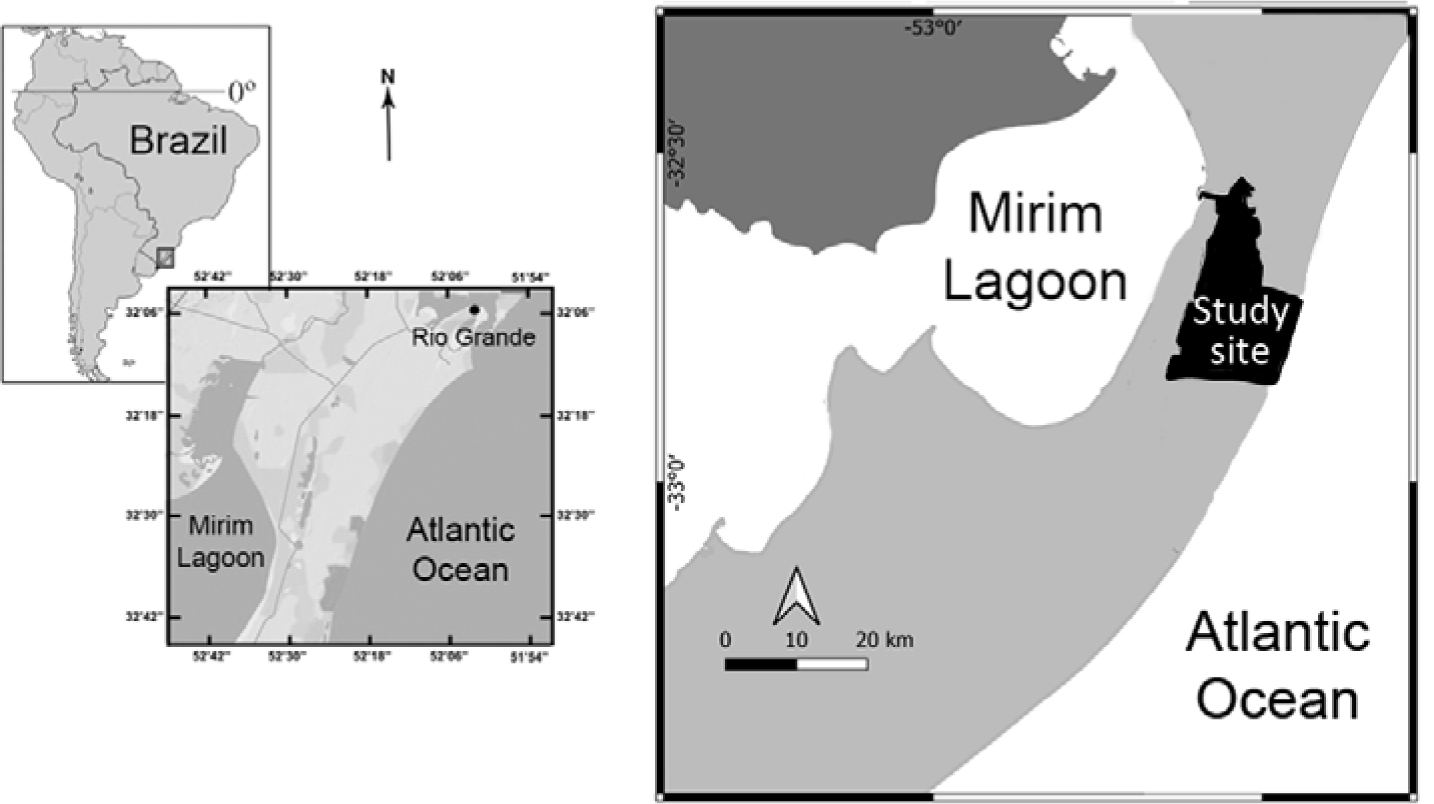
Geographic location of the study area (Estação Ecológica do Taim) in the state of Rio Grande do Sul, Brazil.

The climate of the region is humid sub-temperate, with an annual average temperature of 18.1° C and average temperature of the coldest month of 12.7° C. The annual rainfall is 1162 mm, with periods of drought in the spring and a higher incidence of rain in the winter (Maluf 2000). The maximum temperature varied between 9.1° C and 39.3° C during the study period (May 2011 to April 2012), with the hottest months being February and March 2012; the minimum temperature varied between 0.9 ° C and 28.1 ° C, with the coldest months being July and August 2011. Meteorological data were obtained from the database provided by the Meteorological Station of the Universidade Federal do Rio Grande (FURG), located in the municipality of Rio Grande.

### Anuran sampling

Sampling was carried out in the two most remarkable phytophysiognomies of ESEC Taim: (a) sites of grassland dominance and (b) sites of forest dominance (Figure 2). We were able to distinguish the boundary between these two environments precisely even by a visual inspection. This was possible because the forests are distributed in patches merged along the grassland with well-defined boundaries.

**Figure 2:**
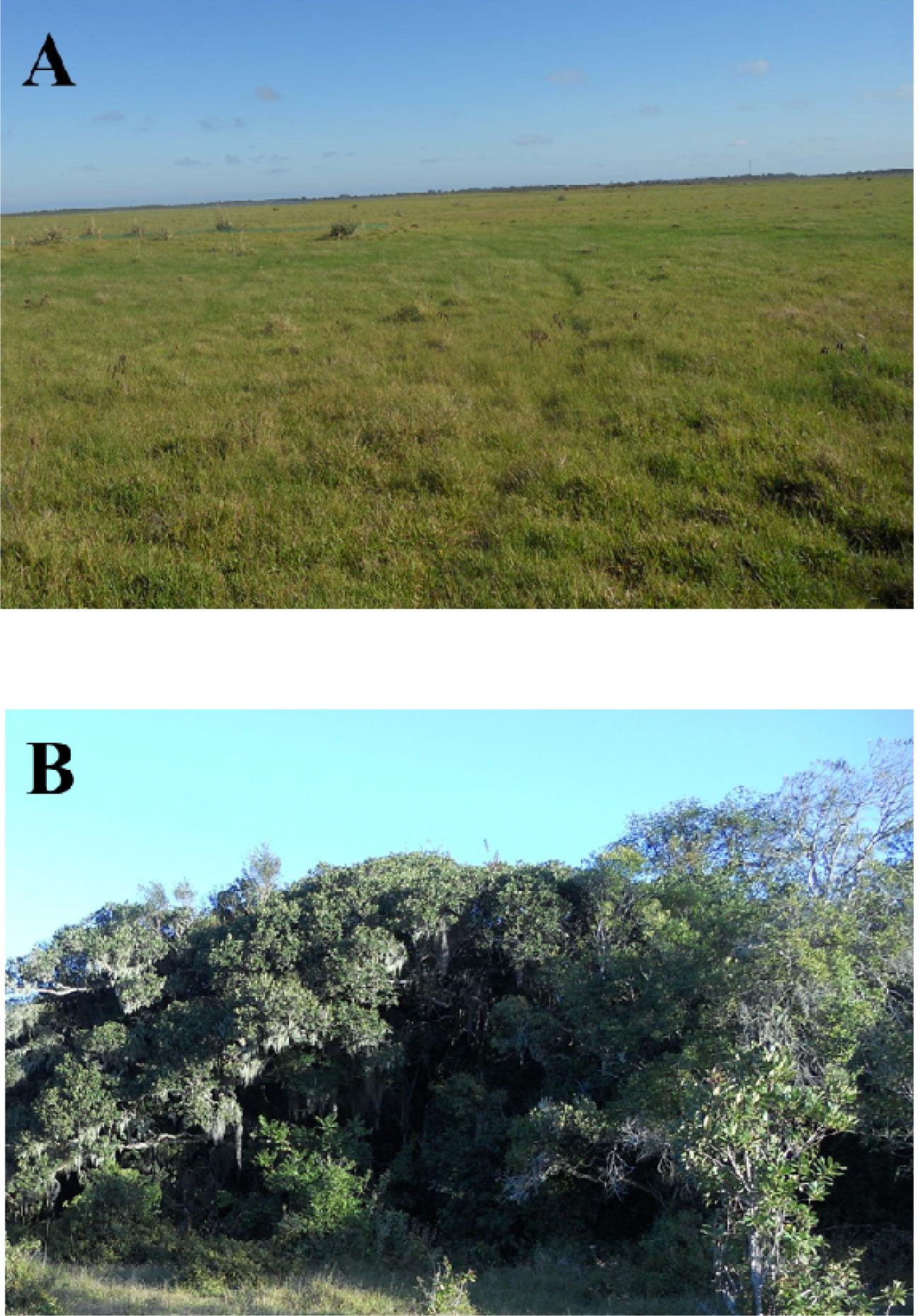
Phytophysiognomies of ESESC Taim: (a) sites of grassland dominance and (b) sites of forest dominance.

We selected two sampling areas in sites of grassland dominance and two in sites of forest dominance. Sampling areas were chosen based on vegetation integrity, area with continuous habitat and presence of potential anuran breeding sites. In addition, we considered our accessibility to these areas since many regions of ESEC Taim are permanently flooded, making the displacement of researchers and the installation of pitfall traps not possible. Sampling areas are at least about 800 m from each other. Field campaigns were carried out every two weeks between May 2011 and April 2012. Each campaign comprised four days when all sites were simultaneously sampled. Frogs were sampled by the combination of three methods: (i) pitfall traps with drift fence; (ii) auditory surveys; (iii) visual search.

### Pitfall traps with drift fence (passive sampling)

We installed sets of buckets arranged in pitfall lines in all the areas of grassland dominance and forest dominance. We installed two lines of buckets in each area, resulting in eight lines and 32 buckets. Pitfalls were arranged following the description of similar studies (e.g. Cechin & Martins 2000). Each pitfall was formed by four 100-l buckets buried about 13.3 m from each other and connected by a mosquito net (60 cm in height) that served as a guide fence. The guide fence was buried 10 cm into the ground to avoid individuals to transpose the fence (Cechin & Martins 2000). All buckets had holes at the bottom to prevent them from being flooded by rain.

We grouped the records obtained from each pair of lines of the same sampling point and consider it to be an assemblage of species. Pitfalls remained open during four consecutive days in intervals of 15 days. During the samplings, we carried out daily inspections in the buckets. We identified every captured animal, which was later released about 20 meters from the trap. The collection was carried out with the authorization number 27755-1 from SISBIO.

### Visual search and auditory surveys (active sampling)

Visual search and auditory surveys (Heyer et al. 1994) were performed during the night, starting one hour after sunset. Active sampling was conducted in the vicinity of the pitfall traps. For standardization of sampling effort, we conducted a visual search in an area of about 1 ha around each pitfall line. Based on our previous experience in auditory surveys, we were able to perform a safe species identification of calling males in a maximum area of 5 ha. This was the covered area for calling surveys at each pitfall line. Active sampling was made along four sampling nights and repeated monthly. Sampling started each night in a different environment (grassland or forest), alternately to avoid the same physiognomy to be always sampled at the same time. We applied two hours per person of active sampling.

### Phytophysiognomy evaluation

To better characterize the phytophysiognomies of the sampling areas, the vegetation structure was measured through plots (following Tozetti & Martins 2008). We characterized the vegetation structure in the vicinity of the areas where pitfall traps were installed. We delimited two plots of 3 m x 10 m, which were distributed perpendicularly in each of the pitfall lines, one of which was placed at the beginning and the other at the end of the line. The following variables of the vegetation were measured in each plot:

### Statistical Analysis

We evaluated differences in the anuran composition in the areas within and between habitat types by using permutational multivariate analysis of variance (PERMANOVA) and multivariate dispersion (BETADISPER). We considered the presence and absence of each specie in each transect. We used the Jaccard index as a dissimilarity measure and performed 9999 permutations. PERMANOVA and BETADISPER were run using the “adonis” and “betadisper” functions in the R package vegan.

**Table 1:** Vegetation descriptors measured in each transect.

| Descriptor | Levels |
| --- | --- |
| Types of land cover | Categories: exposed (without any type of cover); cover by grassy vegetation; presence of shrubs; presence of trees |
| Vegetation cover | Percentual |
| Vegetation height | Centimeters |

## Results

We recorded 13 species of six families in both phytophysiognomies (Figure 3; Table 2). Leptodactylidae and Hylidae were the most representative, with six and four species respectively. The hylid *Hypsiboas pulchellus* occurred exclusively in grasslands and *Scinax squalirostris* occurred exclusively in forests. The number of recorded species varied during the sampling period, with the largest number of species recorded in May (12 species), September and November (11 species) in both phytophysiognomies.

**Table 2:**
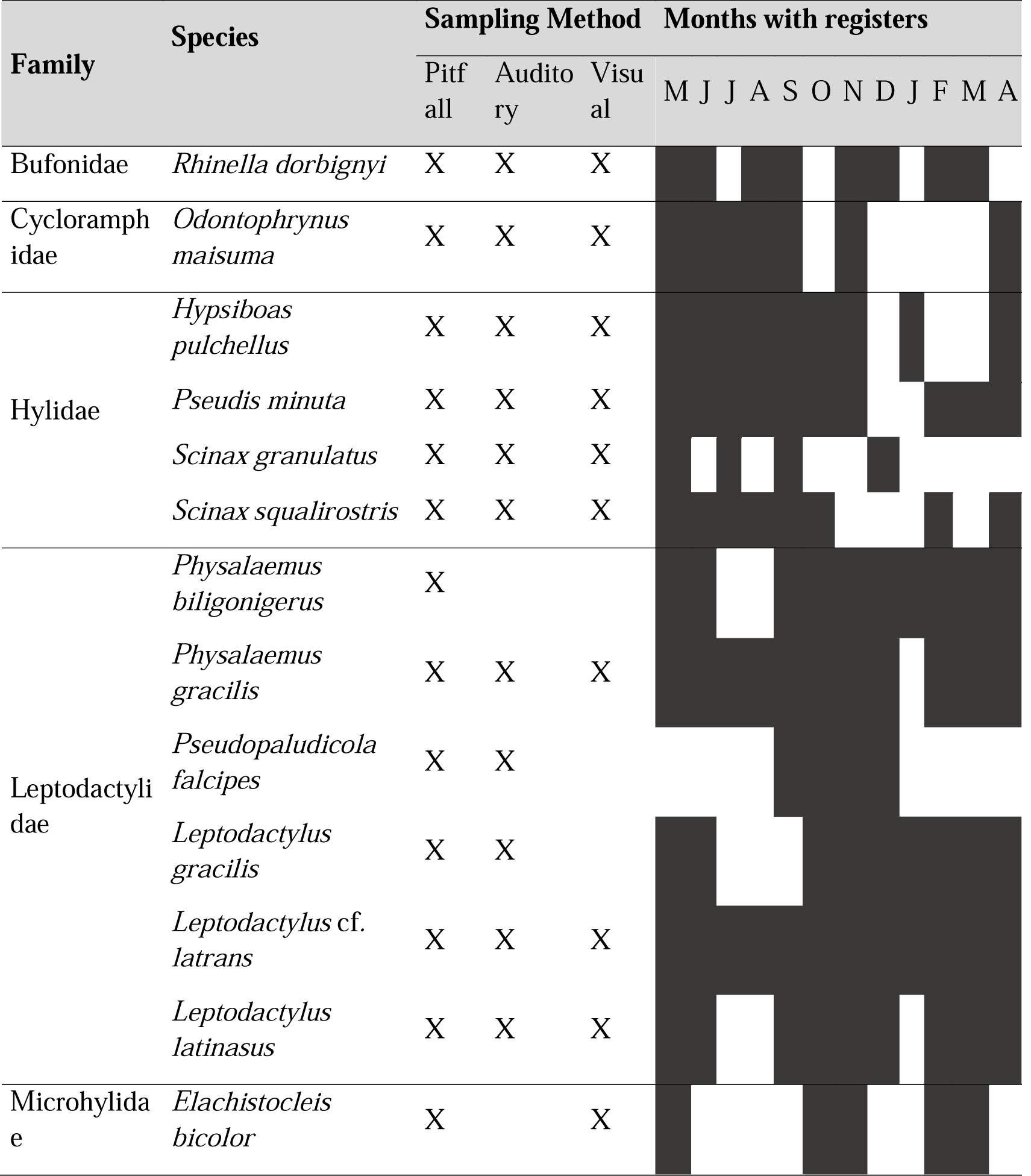
Anuran species found in two phytophysiognomy types of Estação Ecológica do Taim, southern Brazil, between May 2011 and April 2012. Black squares represent the occurrence of species in the relative month.

**Figure 3:**
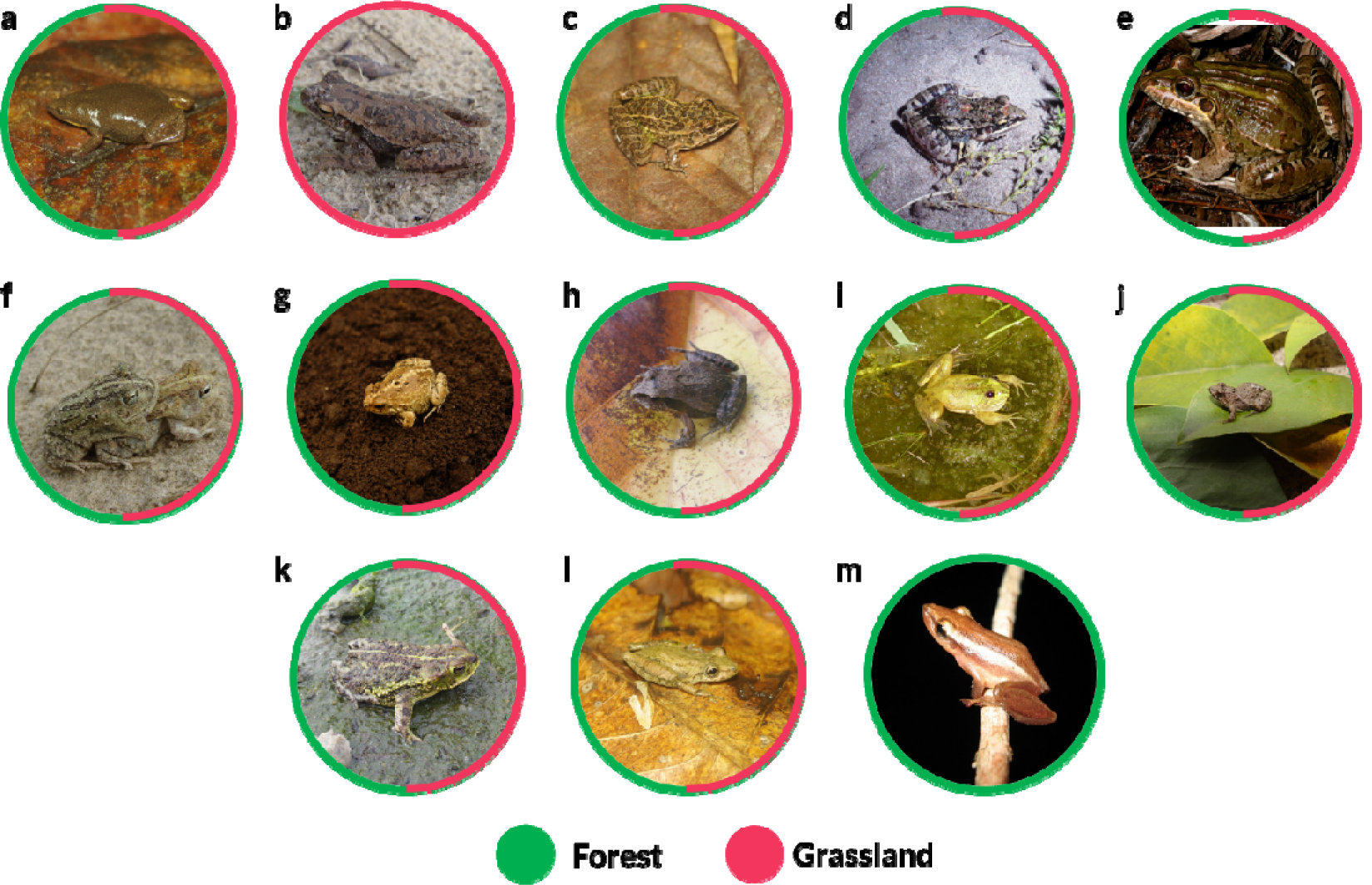
Anuran species found in two phytophysiognomy types of Estação Ecológica do Taim, southern Brazil, between May 2011 and April 2012. (a) *Elachistocleis bicolor*; (b) *Hypsiboas pulchellus*; (c) *Leptodactylus gracilis*; (d) *Leptodactylus latinasus*; (e) *Leptodactylus latrans*; (f) *Odontophrynus maisuma*; (g) *Physalaemus biligonigerus*; (h) *Physalaemus gracilis*; (i) *Pseudis minuta*; (j) *Pseudopaludicola falcipes*; (k) *Rhinella dorbignyi*; (l) *Scinax granulatus*; (m) *Scinax fuscovarius*.

We found that the patterns of compositional dissimilarity (β diversity) were higher between the sampling sites from different phytophysiognomies than within the same phytophysiognomy (*F* = 4.72; *p*=0.05; Figure 4). Also, the difference in composition between the sampling sites in forest areas was considerably high.

**Figure 4:**
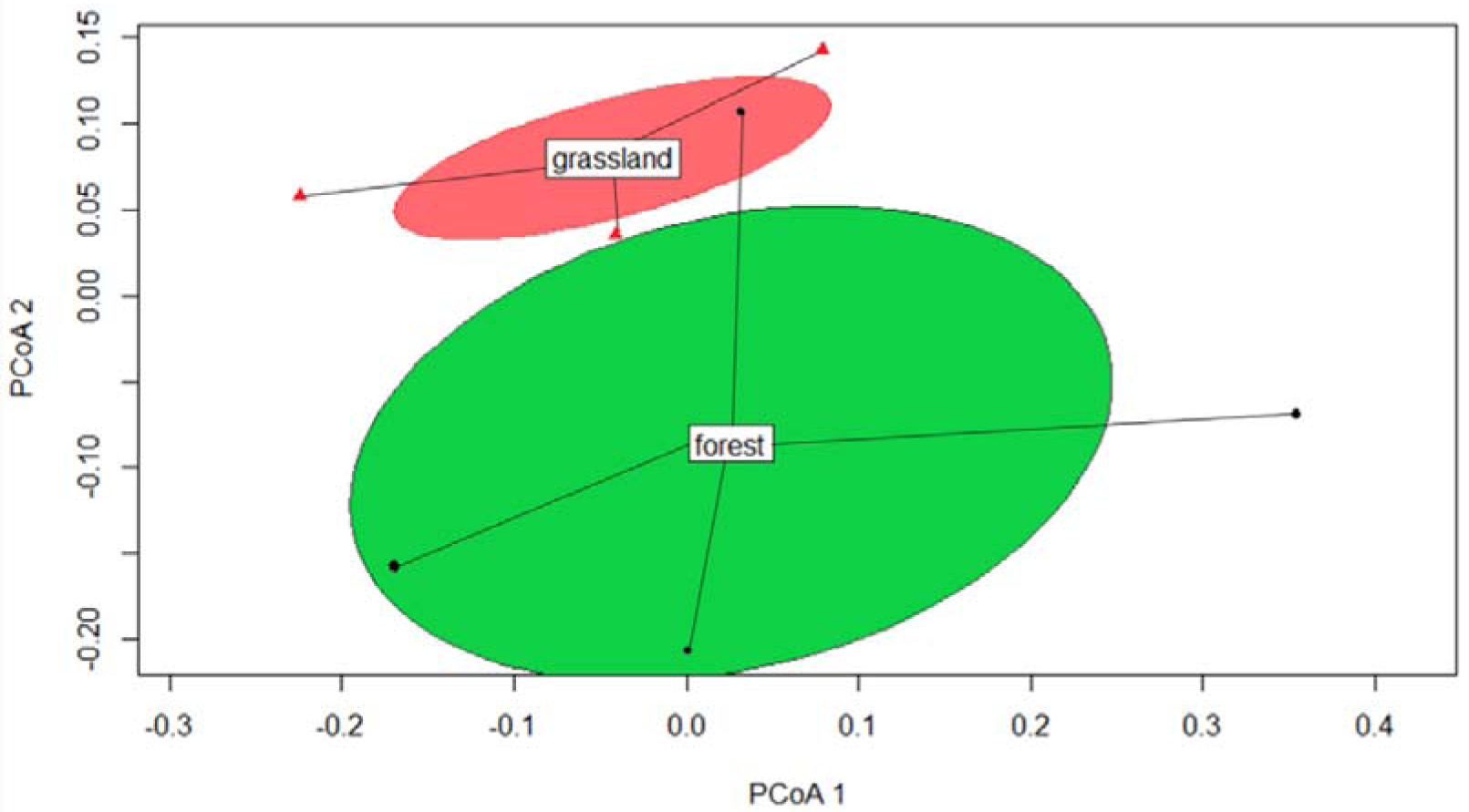
Dissimilarity in anuran composition in two different phytophysiognomies of Reserva Ecológica do Taim, southern Brazil.

## Discussion

Our results showed differences in species composition between grassland and forest habitats. They pointed out that differences in vegetation cover are capable to drive significant changes in anuran species composition. We also highlight that we surveyed adjacent (spatially closely related) sites, which reveals that vegetation has a powerful effect on defining the composition of anuran assemblages. We may assume that the phytophysiognomies evaluated here offer quite different colonization opportunities to anurans, with drastic changes in anuran assemblages across a small geographical area. An obvious relationship between phytophysiognomies and colonization opportunities is related to microhabitat characteristics generated by shading, wind reduction, humidity maintenance, and reduction of dial range of air temperature. Microenvironmental conditions (especially those related to the microclimate) vary substantially between forests and grassland (D’Odorico et al. 2013) and, in our sampling site, mainly because the transition between them is relatively abrupt. Based on these differences, the heterogeneity affects a set of microhabitat proprieties, which could enable the maintenance of some species. The effects of substrate characteristics (e.g., litter deep, moisture retention), and dial variation in air temperature are well-known as determining factors for structuring anuran assemblages (Da Silva et al. 2011b; Van Dyke et al., 2017). Based on this, we point out that the relatively great difference in anuran species composition between sampling sites in forest is a result of the greater heterogeneity of this habitat in relation to grassland. It makes sense since forest sites would vary in both vertical and horizontal components of vegetation cover whereas grasslands vary mostly across the horizontal component (Dalmolin et al. 2019; Dalmolin et al. 2020).

In general, it is expected that forest environments will have a greater variety of microhabitats (Afonso & Eterovick 2007; Prado & Rossa-Feres 2014; Van Dyke et al. 2017). As a result, the occurrence of a greater number of species with different functional traits and evolutionary histories is enhanced (Dalmolin et al. 2019). This possibility opens a precedent for new studies with the specific objective of evaluating the way species use these habitats, as well as their preferences in terms of microhabitats.

We must highlight the fact that *Scinax squalirostris* was recorded only in forests. This result is different from what was expected since this species is associated with low vegetation such as grass and shrubs (Ximenez et al. 2014; Maneyro et al. 2017; Dias et al. 2019). Another aspect to be taken into account is that hylids are less likely to be caught in pitfall traps because of their climbing habits (Oliveira et al. 2013; Dosso et al. 2019). This could have caused some sampling bias and, consequently, affected the observed patterns (which was not our case, since we used different, complementary sampling methods). In addition, some studies indicate that even specific sampling methods such as calling surveys are prone to the observer’s effect on the pattern of results (Dosso et al. 2019).

The fact that we have associated different sampling methods favored the record of species that make wide use of each habitat. Estimations based exclusively on auditory surveys, for example, prioritize the record of habitat use related to reproductive activity (Santos et al. 2016, Moser et al. 2019) limiting the projections regarding habitat requirements (Oliveira et al. 2016; Santos et al. 2016). This is not the case in our study, as previous work carried out in the same region reported the occurrence of nine to fifteen species (Ximenez et al. 2014; Ximenez & Tozetti 2014; Dalmolin et al. 2019; Dalmolin et al. 2020). Thus, our samplings can be considered effective for the representativeness of anuran species in Brazil’s extreme south.

The results herein obtained draw attention to the role of heterogeneity and environmental complexity on anuran composition. They also warn about the imminent risk of changing landscapes due to the invasion of forest species such as *Eucalyptus* and *Pinus* species. Although the sampled communities do not include any endangered species, it is worth mentioning that the study area is a Ramsar site, which is of particular concern when dealing with an area of enormous ecological relevance and importance for the maintenance of many ecosystem functions.

